# The multi-Siglec inhibitor AL009 reprograms suppressive macrophages and activates innate and adaptive tumor immunity

**DOI:** 10.1101/2023.08.02.551627

**Authors:** Sam C. Nalle, Helen Lam, Ling Leung, Spencer Liang, Daniel Maslyar, Arnon Rosenthal

## Abstract

Sialic acid–binding immunoglobulin-type lectins (Siglecs) are cell surface receptors that regulate innate and adaptive immunity, with inhibitory Siglecs promoting immune tolerance. In the tumor microenvironment, overexpression of sialic acid glycans exploits inhibitory Siglec signaling, leading to a cancer-permissive phenotype. AL009 is an engineered Siglec-9-Fc fusion molecule that functions as a sialic acid trap and reprograms suppressive macrophages to activate an anti-tumor immune response. AL009 treatment of human myeloid-derived suppressor cells, an in vitro model of tumor-associated macrophages, resulted in an increase in proinflammatory cytokines and chemokines, changes in cell surface markers, and potent relief of T cell inhibition in a co-culture assay. In syngeneic mouse (*Mus musculus*) tumor models, AL009 engineered with a mouse Fc (AL009m) reduced tumor growth as a monotherapy and in combination with the checkpoint inhibitor anti-PD-L1. In addition, AL009m synergized with the tumor-targeting therapy anti-TRP1 to reduce lung nodules in the B16-F10 intravenous model. Pharmacodynamic marker analysis in syngeneic and humanized mouse tumor models supported an AL009 mechanism of action based on reprogramming tumor-associated macrophages and enhanced T cell activation. Future clinical studies are warranted to further elucidate the safety and efficacy of AL009.

## INTRODUCTION

Sialic acid–binding immunoglobulin-like lectins (Siglecs) are immune regulatory molecules that interact with sialic acid residues in the glycocalyx to modulate innate and adaptive immune response (1–3). Inhibitory Siglecs, such as Siglec-3, -5, -7, -9, and -10, contain an immunoreceptor tyrosine-based inhibition motif that recruits tyrosine phosphatases (such as SHP1 and SHP2) involved in dampening the immune response to maintain homeostasis (1, 2, 4, 5). This is particularly important for recognition of self and resolution of inflammation (6). However, some tumors have exploited this process by overexpressing sialic acid ligands on the tumor cell surface to engage inhibitory Siglecs and evade immune detection, thus promoting an immunosuppressive, cancer-permissive phenotype (1, 5, 7, 8). Disrupting the signaling between immune-inhibitory Siglecs and their sialic acid ligands on hypersialylated cancer cells could confer a therapeutic benefit, making both Siglecs and their sialic acid ligands attractive targets for cancer immunotherapies (2, 8, 9).

Siglecs are expressed on various immune cells but predominantly on myeloid cells, including macrophages and myeloid-derived suppressor cells (MDSCs) (2, 5). The immunoregulatory activity of macrophages, whether proinflammatory or immunosuppressive, can change depending on the chemokines, cytokines, and enzymes in the surrounding environment, with tumor-associated macrophages (TAMs) being primarily immunosuppressive due to the nature of the tumor microenvironment (TME) (5, 10–13). TAMs are not only influenced by the TME but also contribute to the immunosuppressive signaling therein by releasing anti-inflammatory chemokines and cytokines, such as IL-10 and TGF-β, which ultimately lead to T cell inactivation (10, 13). Cell surface markers can be used to monitor the current immunomodulatory status of macrophages; MHC-II, CD80, and CD86 are increased on proinflammatory macrophages, while suppressive macrophages display elevated CD163 and CD206 (5, 13–15). TAM infiltration into tumors has been associated with worse prognosis, particularly when high levels of Siglecs are displayed (16, 17).

Targeted therapeutics designed to reprogram TAMs from an immunosuppressive to a proinflammatory phenotype are currently being investigated (11). Thus far, antibodies against macrophage receptor with collagenous structure (MARCO), the leukocyte immunoglobulin-like receptor B (LILRB) family, Clever-1, and P-selectin glycoprotein ligand-1 (PSGL-1), as well as CD40 agonists, have all shown potential in converting TAMs into proinflammatory effectors (18–22). Interferon gamma (IFNɣ) has also shown potential to skew macrophages towards a proinflammatory phenotype (15, 18, 23). The immunosuppressive characteristic of inhibitory Siglecs along with their expression on TAMs make them ideal targets to modulate macrophage function. However, since multiple inhibitory Siglecs are expressed on TAMs, broad targeting is required to effectively interrupt immunosuppressive Siglec signaling (4, 5).

Here, we present data on AL009, an engineered multi-Siglec inhibitor that functions as a sialic acid trap and reprograms suppressive macrophages to activate an anti-tumor immune response. In vitro, AL009 effectively reprograms suppressive macrophages, as demonstrated by changes in cell surface markers as well as increased secretion of proinflammatory cytokines. In animal tumor models, monotherapy with AL009 or combination therapy with AL009 and anti-PD-L1 or anti-TRP1 reduced tumor burden. Taken together, these data support further investigation of AL009 as a targeted immunotherapy across a broad range of tumors high in Siglec-expressing TAMs.

## MATERIALS AND METHODS

### Study design

The objective of this study was to evaluate the activity of AL009 in vitro and in mouse tumor models. Experiments were designed to elucidate the role of AL009’s Siglec-9 extracellular domain (ECD) and its engineered fragment crystallizable (Fc) region, as well as its interactions with and impact on human myeloid cells. Additionally, AL009 pharmacodynamics and efficacy were tested in mouse models.

### Animals and reagents

C57BL/6 mice harboring a BAC-expressing human Siglec-3, -7, and -9 were generated and maintained by Alector. Expression of human Siglecs was confirmed by flow cytometry. Mice were maintained by crossing Tg^het^ mice with wildtype C57BL/6 mice, and all mice used in studies were Tg^het^ mice. Anti-PD-L1 was produced at Alector based on a reported sequence. Siglec Fcs and isotype control antibodies were produced at Alector unless otherwise stated. In mouse syngeneic tumor studies, a chimeric form of AL009 was produced with a mouse IgG2a Fc (AL009m). huNOG-EXL humanized mice were purchased from Taconic Biosciences (Germantown, NY) and housed at Alector.

### Cell binding competition assay with AL009

MDSCs were generated from healthy human donors by differentiation of CD14^+^ monocytes isolated using a RosetteSep human monocyte enrichment kit (STEMCELL Technologies). In order to generate MDSCs, CD14^+^ monocytes were cultured for 7 days in Roswell Park Memorial Institute media containing 10-ng/mL human granulocyte-macrophage colony-stimulating factor and 10-ng/mL human IL-6 (PeproTech). For cell binding competition with AL009, MDSCs were incubated with titrating amounts of AL009, followed by incubation with Siglec-3, -5, -7, -9, or -10 mouse IgG1 Fc fusion molecules. Bound Siglec-3, -5, -7, -9, and -10 Fcs were assessed by detecting with an anti-mouse IgG-specific secondary (Thermo Fisher) and analyzed via flow cytometry (BD Biosciences).

### Glycan array

We assessed binding to a library of 300 glycan structures for Fc fusion proteins sourced from a commercial vendor (R&D Systems) composed of a fusion of Siglec-3, -5, -7, -9, -10, or -15 ECDs with a human IgG1 Fc. Siglec fusion proteins and control human IgG1 were assessed for glycan binding on the Glycan-300 (RayBiotech). The staining and analysis were performed as a service by RayBiotech using the manufacturer’s protocol. Briefly, the recombinant proteins were incubated with the glycan array for 2 to 3 hours, followed by detection with anti-hIgG biotin and streptavidin-Cy3. Samples were read and fluorescence was normalized across samples. For blocking experiments, AL009 and mIgG1 or mIgG1 alone were applied first before incubation with Siglec Fcs and human IgG.

### MDSC stimulation and RNAseq analysis of MDSCs treated with AL009

MDSCs were treated with AL009 for 48 hours at 37° C and cell surface markers and cytokine production were analyzed via flow cytometry and LEGENDplex (Biolegend), respectively. For RNAseq analysis, MDSCs were treated with 10 µg/mL of AL009 for 24 hours. Cells were harvested and pelleted, followed by resuspension in RLT buffer (Qiagen). RNA was extracted with the RNeasy Plus Microkit (Qiagen). Illumina compatible libraries were created using Lexogen QuantSeq 3’ mRNA-Seq Library Prep Kit FWD and sequenced on a Nextseq P2.

### MDSC**–**T cell co-culture

The activity of AL009 was evaluated alongside several different functional Siglec antibodies in a MDSC–T cell co-culture assay. MDSCs were harvested and co-cultured with autologous CD8^+^ T cells in the presence of Dynabeads Human T-activator CD3/CD28 (Thermo) and either a Siglec-Fc fusion or antibodies. Proliferation via carboxyfluorescein succinimidyl ester (Thermo Fisher) dilution and cytokine production via LEGENDplex were assessed after 3 to 5 days.

### MDSC stimulation with plate-bound IgG and anti-TREM1/anti-TREM2

MDSCs were stimulated with plate-bound human IgG1 with or without 10-µg/mL Siglec-9 ECD co-incubation for 48 hours followed by analysis of TNF production (Meso Scale Discovery). Separately, MDSCs were stimulated with anti–triggering receptor expressed on myeloid cells 1 (TREM1) or anti–triggering receptor expressed on myeloid cells 2 (TREM2) agonistic antibodies with or without Siglec-9 ECD for 48 hours followed by analysis of TNF production.

### Fcγ receptor blockade

MDSCs were incubated with the 10-µg/mL FcγRIIa (clone IV.3)- or FcγRIIb (clone 2B6)-blocking antibodies alone or in combination for 30 minutes on ice, followed by treatment with 3.3-µg/mL AL009 for 24 hours at 37° C and analysis of cell surface markers by flow cytometry.

### Specific binding of AL009

For peripheral immune cell binding specificity, healthy human blood was incubated with AF647-labeled AL009 using SiteClick reagents (Thermo Fisher) followed by analysis via flow cytometry. For myeloid binding, healthy human monocytes from peripheral blood, M0 macrophages differentiated from human monocytes with 50-ng/mL human M-CSF, M1 macrophages differentiated from human monocytes with 50-ng/mL M-CSF, 100-ng/mL lipopolysaccharide (InvivoGen), and 50-ng/mL IFNγ (PeproTech), or MDSCs, were incubated with AF647-labeled AL009 followed by analysis via flow cytometry.

### Biodistribution of AL009

Female C57BL/6 mice bearing established subcutaneous MC38.NCI.TD1 murine colon carcinoma were injected intravenously with 10-mg/kg AF647-labeled AL009m. Ex vivo fluorescence imaging using the IVIS Spectrum imaging system was performed on liver, spleen, heart, and lungs collected from each mouse at 1, 6, 24, 48, and 96 hours post injection. Three animals were imaged per time point.

### Effect of AL009 in mouse models

The MC38 and B16-F10 cell lines were purchased from American Type Culture Collection. The E0771 cell line was purchased from MI Bioresearch. Female S3/7/9 Tg mice that were 7 to 12 weeks old were implanted with MC38 or E0771 cells subcutaneously in the flank region at a density of 1.0 × 10^5^ or 4.0 × 10^5^ cells per mouse, depending on the model. When tumors reached an average volume of ∼100 mm^3^, mice were randomized into treatment groups and dosing was initiated. Mice received intraperitoneal injections of 10- to 20-mg/kg AL009m or isotype control with or without 3-mg/kg anti-PD-L1 twice per week for 3 weeks. Tumors were measured twice per week with calipers. For the B16-F10 intravenous model, 1.5 × 10^5^ B16-F10 cells were administered via intravenous injection. Mice were treated with 27-µg/mouse anti-TRP1 (BioXCell) on day 1 in addition to 10-mg/kg AL009m or isotype control on days 1, 4, 7, 10, and 13. A satellite group of animals was dosed only with control antibodies, and several mice in this group were euthanized every 2 days starting on day 9 to assess lung nodules. When the satellite group reached 50 to 100 lung nodules, the remaining experimental groups were euthanized for analysis. Statistical testing was performed by 2-way Student’s t-test.

### Pharmacodynamic biomarkers

Siglec-3/7/9 BAC Tg mice were implanted with E0771 tumors and randomized for treatment as described above. The mice received intraperitoneal 20-mg/kg AL009m or isotype control on days 0, 4, and 7, and were sacrificed on day 8. Splenic and tumor tissues were mechanically dissociated and passed through a 70-μm filter to obtain single-cell suspensions. Peripheral blood was treated with ammonium-chloride-potassium lysis buffer to lyse red blood cells and washed with phosphate-buffered saline. Cell surface markers on immune cell subsets were analyzed by flow cytometry. In order to identify immune cell subsets, CD3+ T cells were plotted on the y-axis and CD11b+ myeloid cells were plotted on the x-axis, with further subgating occurring from there.

For pharmacodynamic analysis of humanized mice, 14- to 16-week-old female huNOG-EXL mice were implanted with A375 melanoma cells subcutaneously in the flank region at a density of with 3.0 × 10^6^ cells per mouse. When tumors reached an average volume of 200 to 400 mm^3^, mice received intraperitoneal injections of 10-mg/kg AL009 or isotype control on days 0 and 3 and were sacrificed 24 hours later. Cell surface markers on immune cells subsets were analyzed by flow cytometry.

### Statistical analyses

Statistical analysis was performed using GraphPad Prism v9. Data are displayed as mean ± SEM. A 2-tailed t-test was used for comparisons between 2 groups. A 1-way ANOVA followed by the Holm-Sidak method of multiple comparisons was used for comparisons of more than 2 groups. A *p* value of < 0.05 was considered significant.

### Study approval

All in-house animal studies were reviewed by, approved by, and performed in compliance with Alector’s Institutional Animal Care and Use Committee protocols. Outsourced studies were conducted in accordance with the applicable Covance or Charles River Laboratories standard operating procedures, which approved these studies.

## RESULTS

### AL009 displays specific blocking between inhibitory Siglecs and their ligands

AL009 is composed of a Siglec-9 ECD and an engineered Fc to localize the therapeutic to immune cells and promote cooperative binding of the ECD. Given the redundancy of inhibitory Siglecs, an effective therapeutic would require broad blocking between multiple inhibitory Siglecs and their respective ligands. To assess the degree to which AL009 could disrupt Siglec-ligand binding on MDSCs, the binding between Siglec-3, -5, -7, -9, and -10 Fc fusion proteins and human MDSCs was quantified in the presence of increasing concentrations of AL009. With all the Siglec Fcs tested, an AL009 dose-dependent decrease in binding was observed, demonstrating a broadly effective blockade of ligand recognition on cells (Fig. 1A). In order to assess specific ligand blockade, a biochemical blocking assay using AL009 was established with a glycan array containing over 100 sialic acids. Using this assay format, AL009 blocked binding of known sialic acids recognized by inhibitory Siglecs -3, -5, and -7 (Fig. 1B-D). As the binding portion of AL009 is composed of a Siglec-9 ECD, Siglec-9 binding relative to other inhibitory Siglecs was evaluated side by side using a glycan array. Siglec-9 displayed an extensive sialic acid binding profile compared to other inhibitory Siglecs (Fig. S1), helping to explain the ability of AL009 to block ligands recognized by multiple Siglecs on MDSCs. Overall, these results demonstrate that AL009 possesses the ability to block specific ligand binding of at least 5 inhibitory Siglecs present on myeloid cells.

**Fig. 1.**
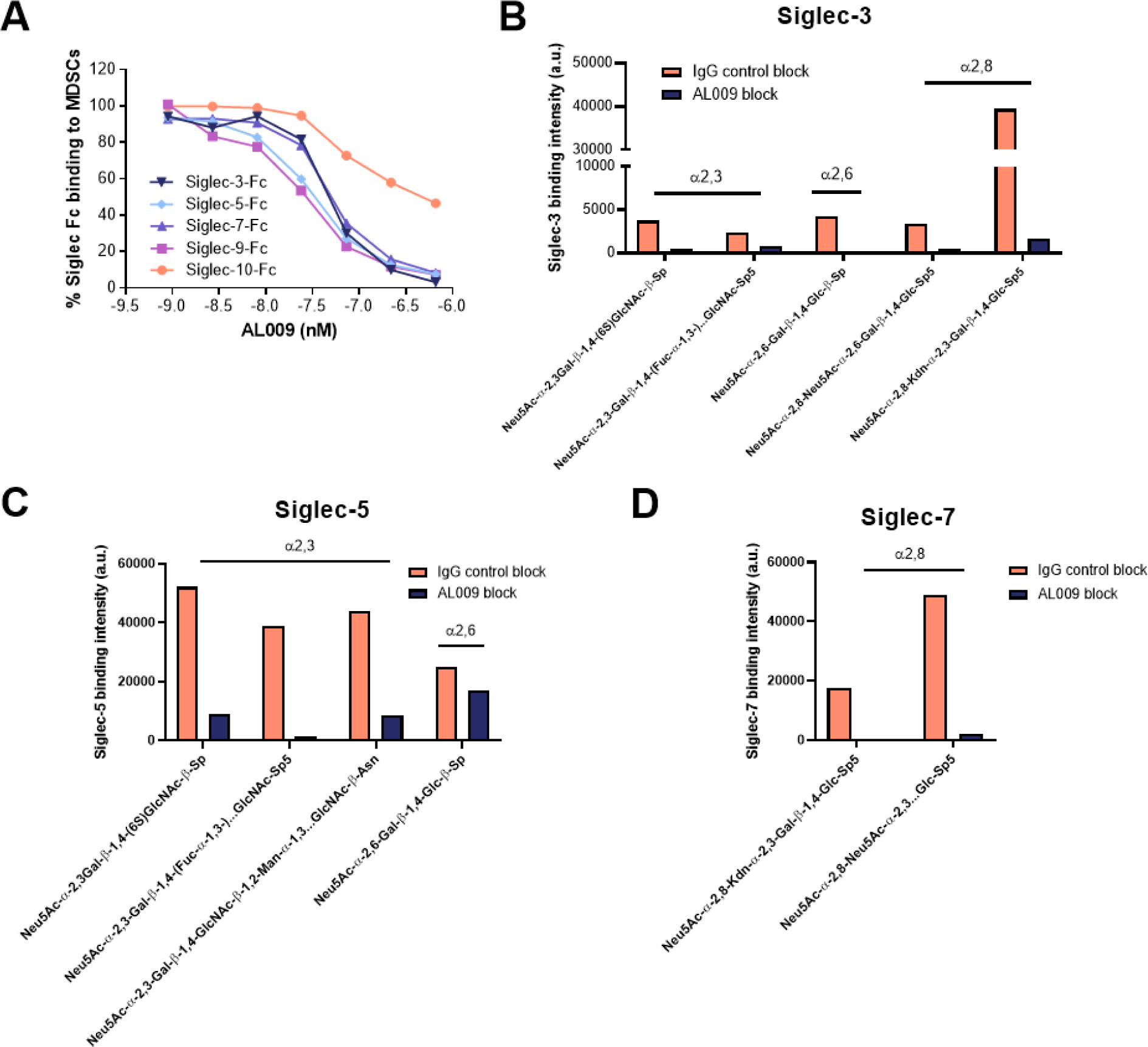
AL009 blocks binding of inhibitory Siglec receptors and their ligands. (**A**) AL009 blocked binding of Siglec-3, -5, -7, -9, and -10 from binding to MDSCs in a dose-dependent manner. (**B-D**) In a glycan array containing over 100 sialic acids, AL009 blocked Siglec-3, -5, and -7 from binding their respective ligands. Fc, fragment crystallizable; MDSCs, myeloid-derived suppressor cells; Siglec, sialic acid–binding immunoglobulin-type lectin.

### AL009 reprograms MDSCs

The phenotypic effect of AL009 blocking recognition between inhibitory Siglecs and their ligands was assessed by incubating MDSCs with AL009 and evaluating cell surface markers, cytokines, and chemokines. AL009 treatment resulted in a dose-dependent increase in immune activation marker CD86 and a dose-dependent decrease in the immunosuppressive macrophage marker CD163, consistent with reprogramming of MDSCs (Fig. 2A, 2B). Further, TNF production was increased in a dose-dependent manner (Fig. 2C). In order to gain a greater understanding of the reprogramming effect of AL009 on MDSCs, RNAseq analysis was performed on MDSCs after AL009 treatment. AL009 induced robust expression of cytokines, chemokines, and co-stimulatory molecules compared to isotype control in MDSCs (Fig. 2D).

**Fig. 2.**
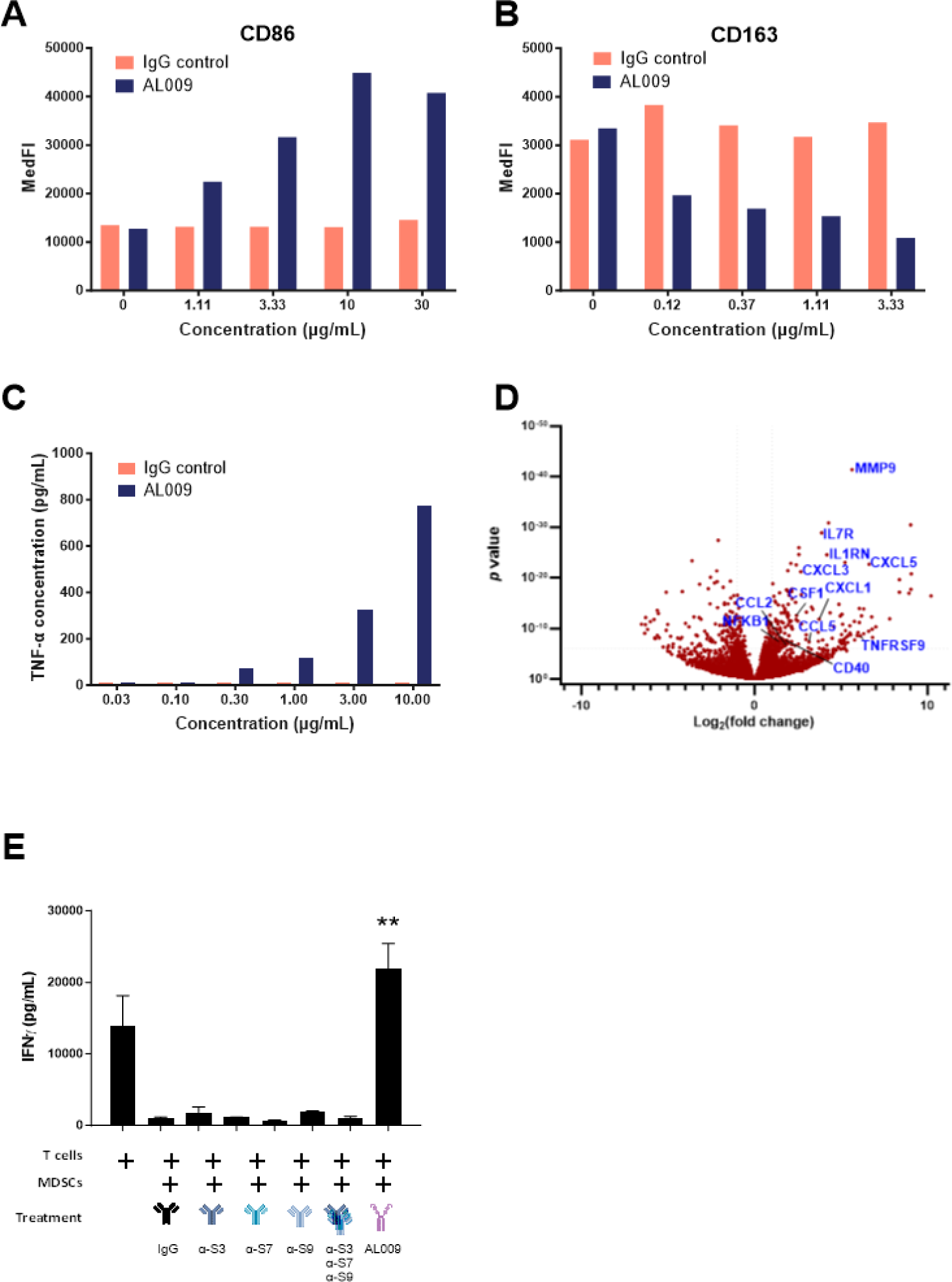
AL009 repolarizes suppressive MDSCs. AL009 (**A**) enhanced CD86, (**B**) repressed CD163, and (**C**) induced TNF secretion after treatment on MDSCs. (**D**) AL009 treatment of MDSCs led to robust changes in gene expression. (**E**) AL009 rescued T cells suppressed by MDSCs in a co-culture system. Error bars represent SEM. IFNɣ, interferon gamma; MDSCs, myeloid-derived suppressor cells; MedFI, median fluorescence intensity; Siglec, sialic acid– binding immunoglobulin-type lectin. \*\**p* < 0.01, 2-sided t-test.

We considered whether the reprogramming effect of AL009 treatment was specific to the Siglec-9 ECD portion of the molecule. To evaluate this, MDSCs were treated with Siglec-3, -5, -7, -9, or -10 ECD-Fc fusions and the reprogramming effect was assessed. The Siglec-9–mIgG fusion resulted in the largest increase in activation markers CD86 and CD40 and the largest reduction in suppressive macrophage markers CD163 and CD206 (Fig. S2A-D). The Siglec-9– mIgG fusion was also the only Siglec-Fc fusion that induced a dose-dependent increase in CCL4 production by MDSCs, consistent with reprogramming of these cells (Fig. S2E).

Next, we examined whether AL009 was able to rescue MDSC suppression of T cells in a co-culture system. MDSCs were treated with ligand-blocking antibodies against Siglec-3, -7, or -9, a combination of all 3 antibodies, or AL009 prior to co-culture with activated T cells. The amount of IFNɣ produced by the T cells was reduced in the presence of MDSCs and was not rescued by any treatment except AL009 (Fig. 2E). In addition to increasing IFNɣ production in T cells, we tested AL009’s impact on T cell proliferation. After treatment with increasing concentrations of AL009, T cell proliferation was quantified, with a half maximal effective concentration of 1 to 2 nM AL009 (Fig. S3). Overall, AL009 is effective at potently repolarizing MDSCs, resulting in phenotypic changes consistent with a proinflammatory response and relief of T cell inhibition.

### AL009 activity depends on the ECD and Fc regions

We hypothesized that both the Siglec-9 ECD and engineered Fc regions of AL009 were important for the function of the molecule. To determine the contribution of the Siglec-9 ECD portion of AL009, we first created domain mutation constructs. Production of the chemokine CCL5 was induced with AL009, but that effect was ablated when the Siglec-9 ECD portion of AL009 was deleted (Fig. 3A). Next, the Siglec-9 ECD and Fc fusion portions of AL009 were decoupled, and MDSCs were incubated with plate-bound IgG to engage Fcγ receptors (FcγRs), with or without a soluble AL009 mutant consisting of only the Siglec-9 ECD, then analyzed for TNF production as a marker of MDSC programming (Fig. 3B). FcγR engagement by increasing concentrations of plate-bound IgG induced a low level of TNF that was enhanced by the addition of Siglec-9 ECD (Fig. 3C). We next asked if the Siglec-9 ECD could augment a separate activation signal beyond FcγRs. Soluble anti-TREM1 or anti-TREM2 agonist antibodies were incubated with MDSCs at increasing concentrations in addition to Siglec-9 ECD. Anti-TREM1 and anti-TREM2 had very minimal effects on MDSCs alone, but the addition of a fixed amount of Siglec-9 ECD resulted in increased secretion of TNF in combination with TREM1 or TREM2 antibodies (Fig. 3D). These results demonstrate that Siglec-9 ECD alone is active and synergizes with an activation signal in order to reprogram MDSCs.

**Fig. 3.**
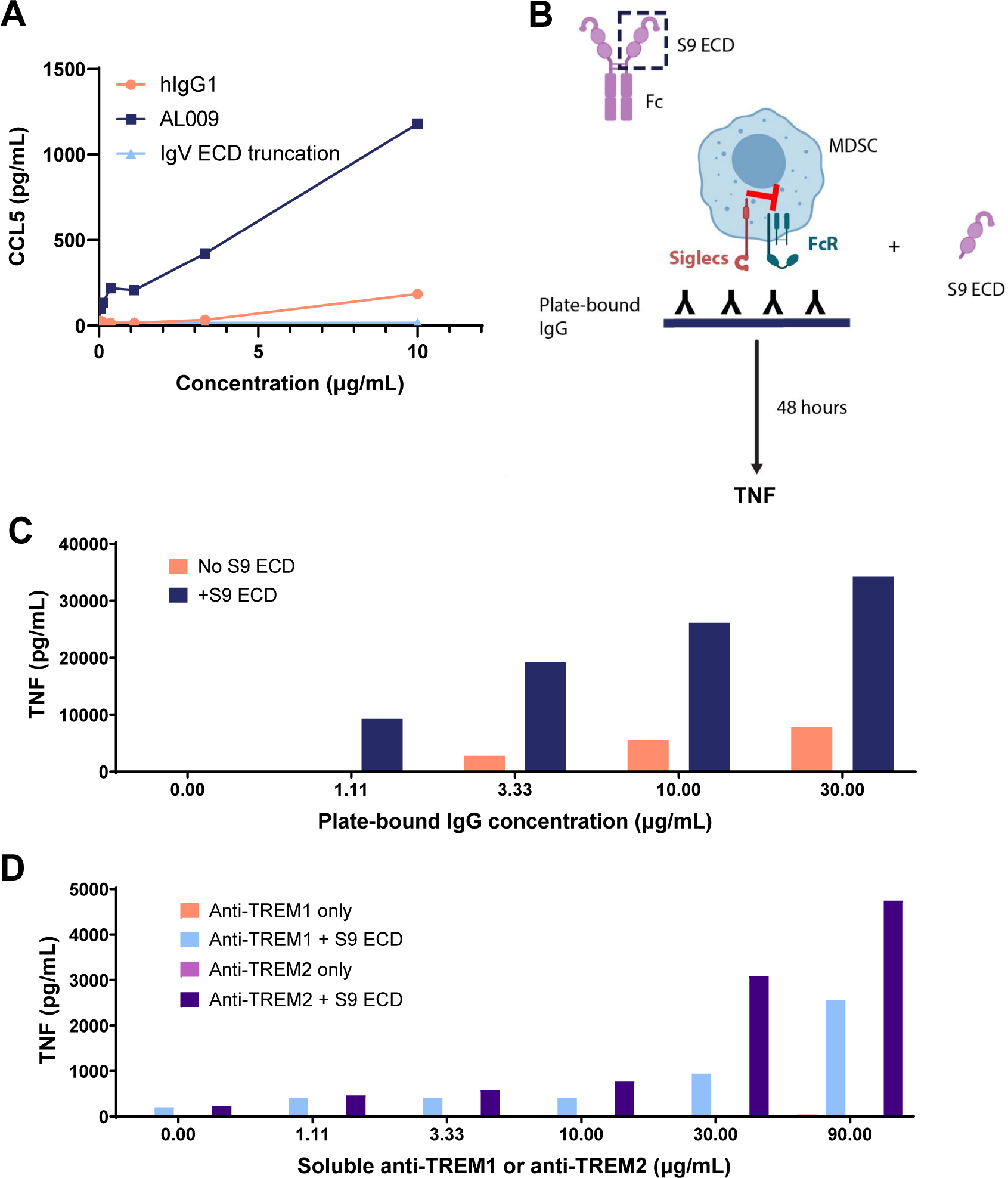
The Siglec-9 ECD relieves immune suppression in the presence of an activating signal. (**A**) Truncation of the ECD domain of AL009 ablated its ability to induce CCL5. (**B**) Schematic of experimental setup. (**C**) Levels of TNF produced by MDSCs upon IgG cross-linking were increased in the presence of Siglec-9 ECD. (**D**) Levels of TNF produced by MDSCs were not induced by anti-TREM1 or anti-TREM2 alone but were induced when Siglec-9 ECD was present. ECD, extracellular domain; Fc, fragment crystallizable; MDSCs, myeloid-derived suppressor cells; MedFI, median fluorescence intensity; Siglecs, sialic acid–binding immunoglobulin-type lectins.

The contribution of the Fc portion of AL009 was also investigated. The ability of AL009 to relieve T cell suppression in an MDSC–T cell co-culture assay was greatly reduced when the Fc portion of AL009 was silenced via mutations that ablate binding to FcγRs (Fig. 4A). To further examine the contribution of the Fc portion of AL009 in binding MDSCs, dissociation constants on cells were determined. The dissociation constant for AL009 when incubated with MDSCs was 1.3 nM. However, when non-Fc receptor–expressing cells were used in place of MDSCs, binding was much weaker (Kd = 220 nM). Similarly, when the Fc portion of AL009 was silenced, binding to MDSCs was greatly reduced (Kd = 97 nM). In order to determine which Fc gamma receptors were important for AL009 function, the ability of AL009 to increase CD86 expression at the cell surface, as a marker of MDSC reprogramming, was assessed in the presence of antibodies against FcɣRIIa, FcɣRIIb, or a combination of FcɣRIIa and FcɣRIIb. The N325S/L328F (NSLF) mutation contained within the Fc portion of AL009 prevents binding to FcɣRIIIa and FcɣRIIIb; therefore, as expected there was no effect when MDSCs were incubated with an FcɣRIII blocking antibody prior to AL009 treatment. FcɣRIIa blockade reduced AL009 activity, while FcɣRIIb had no effect. Consistent with these results, the combination of FcɣRIIa- and FcɣRIIb-blocking antibodies was equivalent to FcɣRIIa blockade alone (Fig. 4B). Taken together, these results suggest FcɣRIIa is necessary for AL009 activity on MDSCs.

**Fig. 4.**
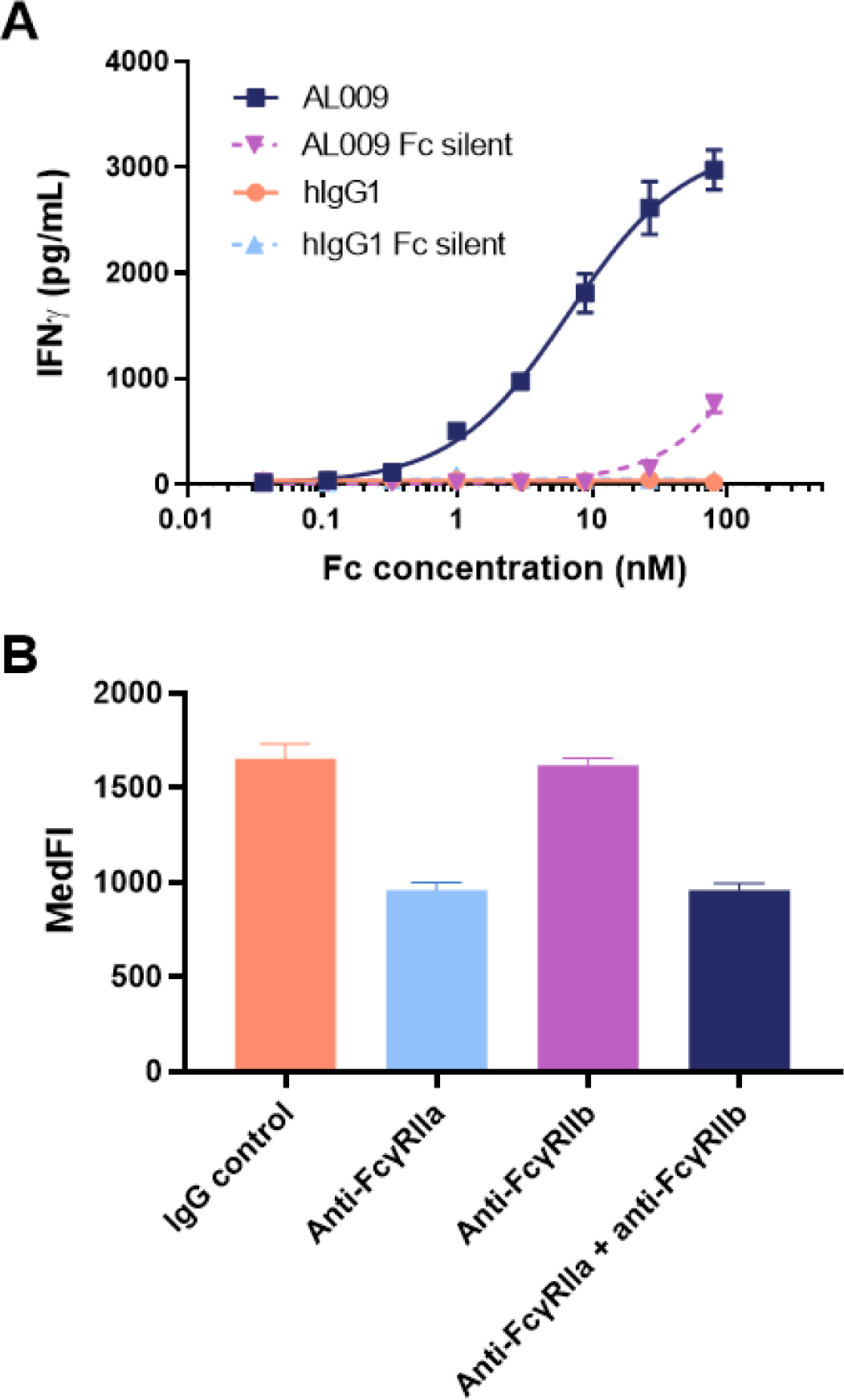
AL009 activity requires Fc receptor interactions. (**A**) While AL009 was able to reverse MDSC suppression and lead to higher levels of IFNɣ compared to control, AL009 Fc silent was not able to elevate IFNɣ to the same extent. (**B**) FcɣRIIa was required for functional activity of AL009 on MDSCs. Data are shown as mean ± SEM. Fc, fragment crystallizable; IFNɣ, interferon gamma; MDSC, myeloid-derived suppressor cell; MedFI, median fluorescence intensity.

### AL009 preferentially binds myeloid cells

In whole human blood, binding of AL009 to a variety of immune cell subsets, including monocytes, granulocytes, B cells, natural killer cells, T cells, and platelets, was assessed. AL009 preferentially bound monocytes compared to the other immune cell subsets tested (Fig. 5A). To further elucidate cell subsets that are preferentially bound by AL009, donor blood samples were used to compare AL009 binding to monocytes, macrophages, and MDSCs. The highest binding observed was between AL009 and MDSCs, followed by non-activated M0 macrophages, and then classically activated M1 macrophages (Fig. 5B).

**Fig. 5.**
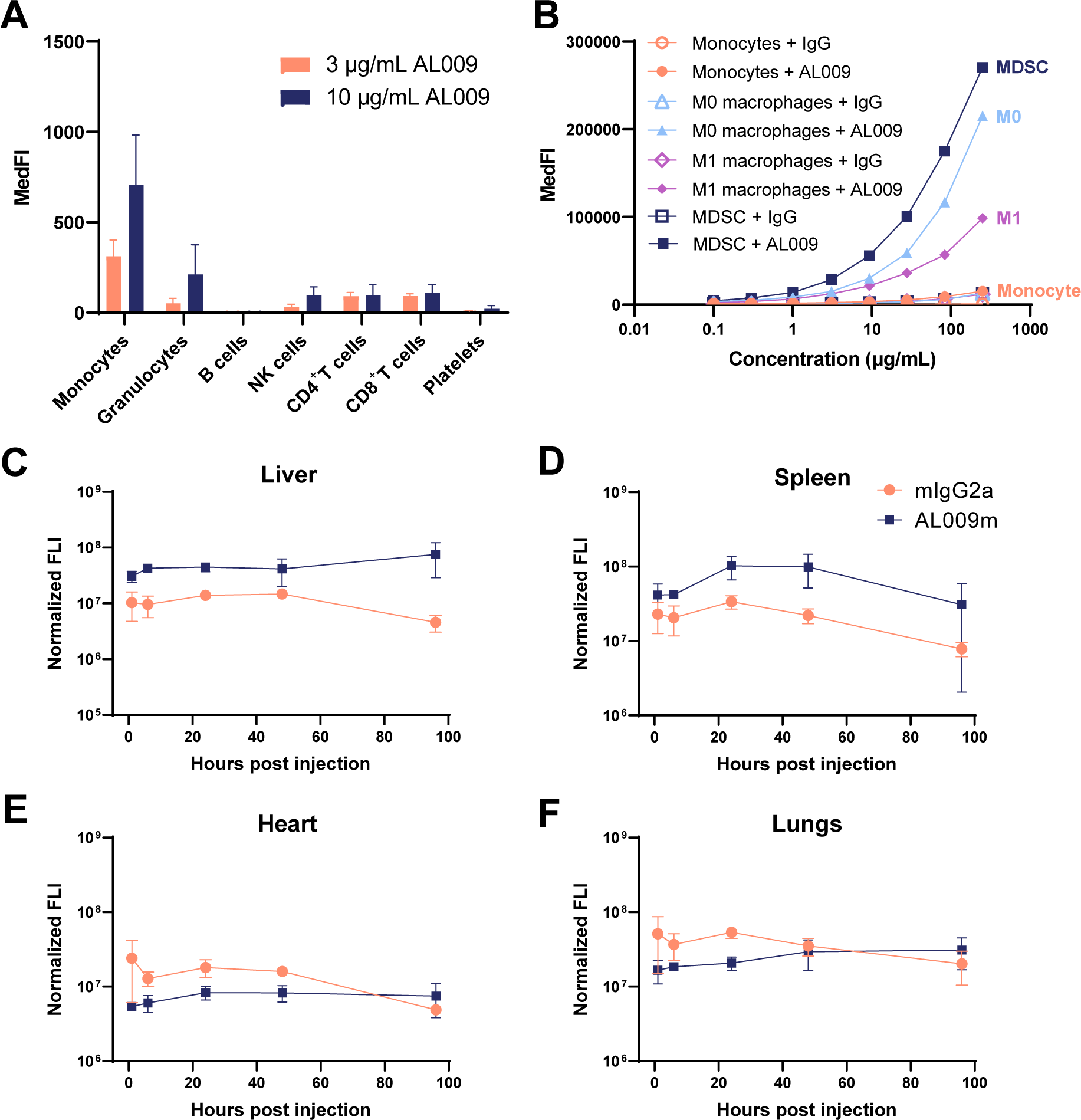
AL009 preferentially binds myeloid cells. (**A**) In whole human blood, AL009 preferentially bound monocytes over granulocytes, B cells, NK cells, T cells, and platelets. (**B**) AL009 preferentially bound MDSCs over other myeloid subsets, including M0 and M1 macrophages and monocytes. (**C-F**) In a biodistribution study in mice, AL009m accumulated in macrophage-rich tissues, such as liver and spleen (n = 3 mice per timepoint). Error bars represent SEM. FLI, fluorescence intensity; MDSC, myeloid-derived suppressor cell; MedFI, median fluorescence intensity; NK, natural killer.

AL009m, a mouse reactive surrogate for AL009 containing a mouse Fc, was created to assess the activity of AL009 in vivo. A biodistribution study was performed with AL009m by fluorescently labeling the molecule and then injecting it intravenously in mice. AL009m accumulation was higher than the isotype control in macrophage-rich mouse tissue samples, such as liver and spleen, while no difference between AL009m or isotype control was seen in other tissues, such as heart and lung (Fig. 5C-5F). The results from both the whole human blood binding study and biodistribution study provide evidence that AL009 preferentially binds myeloid cells both in vitro and in vivo.

### AL009m demonstrates anti-tumor activity in mouse syngeneic tumor models

Several syngeneic mouse tumor models were used to assess the anti-tumor efficacy of AL009m in vivo. As a monotherapy, AL009m was tested in the E0771 triple negative breast cancer model. Tumor volume was reduced and survival improved with AL009m treatment compared to isotype control (Fig. 6A, 6B). To assess the impact of AL009m in combination with other therapies, the B16-F10 intravenous injection model in combination with anti-TRP1 was utilized. In this model, anti-TRP1 induces macrophage-mediated antibody-dependent cellular phagocytosis (ADCP) and antibody-dependent cellular cytotoxicity (ADCC) of B16-F10 melanoma cells (24). Combination treatment with AL009m and anti-TRP1 resulted in a significant reduction of lung nodules compared to anti-TRP1 and isotype control (Fig. 6C). These results demonstrate that AL009m can enhance ADCP/ADCC in vivo. To test the combination of AL009 with checkpoint inhibitors, anti-PD-L1 was utilized along with AL009m in the MC38 and E0771 models. In both models, anti-PD-L1 alone was not significantly different than isotype control, whereas combination treatment with AL009m led to a significant reduction in tumor volume (Fig. 6D, 6E). Taken together, these efficacy results demonstrate that AL009m is active as a monotherapy and in combination with an agent that induces ADCP/ADCC and the representative checkpoint inhibitor anti-PD-L1.

**Fig. 6.**
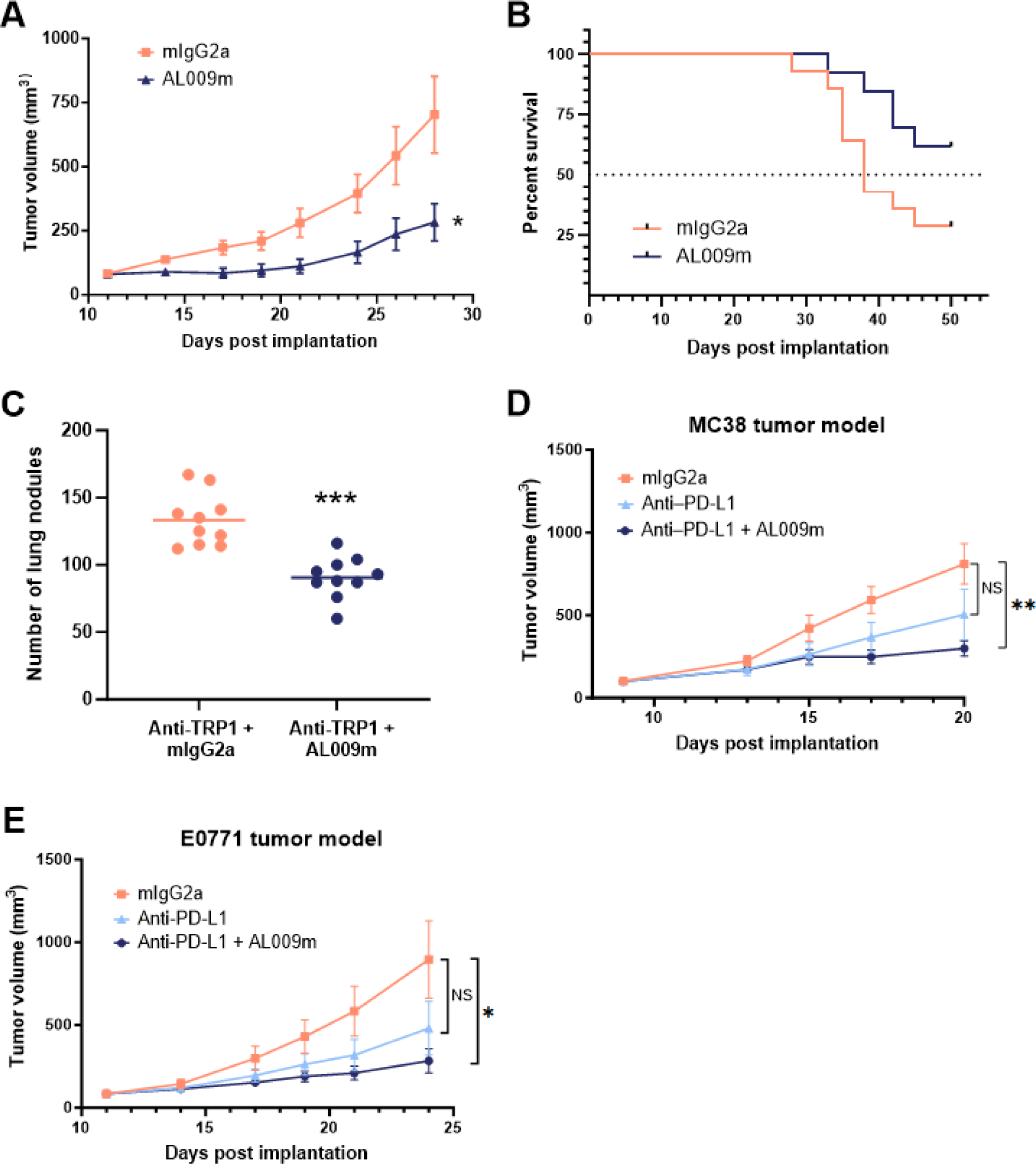
AL009m demonstrated anti-tumor activity in multiple mouse syngeneic tumor models. (**A**) In the E0771 breast cancer model, AL009m treatment resulted in a significant reduction in tumor volume as well as (**B**) increased survival. (**C**) In the intravenous B16-F10 tumor model, combination treatment with anti-TRP1 and AL009 resulted in reduced lung nodules. Line at mean. (**D**) In the MC38 tumor model, combination treatment with AL009m and anti-PD-L1 resulted in reduced tumor volume. (**E**) In the E0771 breast cancer model, combination treatment with AL009m and anti-PD-L1 showed a synergistic effect. (D-E) Data are shown as mean ± SEM. \**p* < 0.05, \*\**p* < 0.01, \*\*\**p* < 0.001, 2-sided t-test.

### Pharmacodynamic markers show enhanced immune activation in the tumor and spleen following AL009 treatment

To evaluate the pharmacodynamic effect of AL009m in the context of a tumor, spleen and tumor samples from the E0771 mouse tumor models were analyzed by flow cytometry following treatment with AL009m alone or in combination with anti-PD-L1. As a monotherapy, AL009m treatment resulted in a significant decrease in Ly6C^+^ cells, as a percentage of CD11b^+^ cells, indicating a decrease in MDSCs in tumor tissue (Fig. 7A). In spleen samples, an increase in HLA-DR with AL009m treatment signaled activation of splenic macrophages (Fig. 7B). Further, following AL009m treatment, there was a concurrent increase in activated CD4^+^ cells in the spleen, as measured by an enhanced percentage of CD69^+^ cells (Fig. 7C). There was also an increase in activated CD8^+^ cells induced by AL009m that was further enhanced by anti-PD-L1 treatment, indicating a synergistic effect (Fig. 7D). In addition to the effects observed in the tumor and spleen, AL009m also induced a dose-dependent increase in surface CD86 on circulating monocytes (Fig. S4), similar to what was observed on human MDSCs following AL009 treatment (Fig. 2A). The pharmacodynamic effect of AL009 was further evaluated in a humanized mouse tumor model. In this humanized model, immunodeficient mice were engrafted with human CD45^+^ stem cells, implanted with a human A375 tumor weeks later, and treated with AL009 approximately 2 weeks post implantation. Flow cytometry was used to analyze tumor and spleen samples. Treatment with AL009 significantly decreased surface CD163 on TAMs and splenic macrophages (Fig. 7E, 7F), demonstrating an in vivo reprogramming effect consistent with the in vitro reprogramming observed with human MDSCs.

**Fig. 7.**
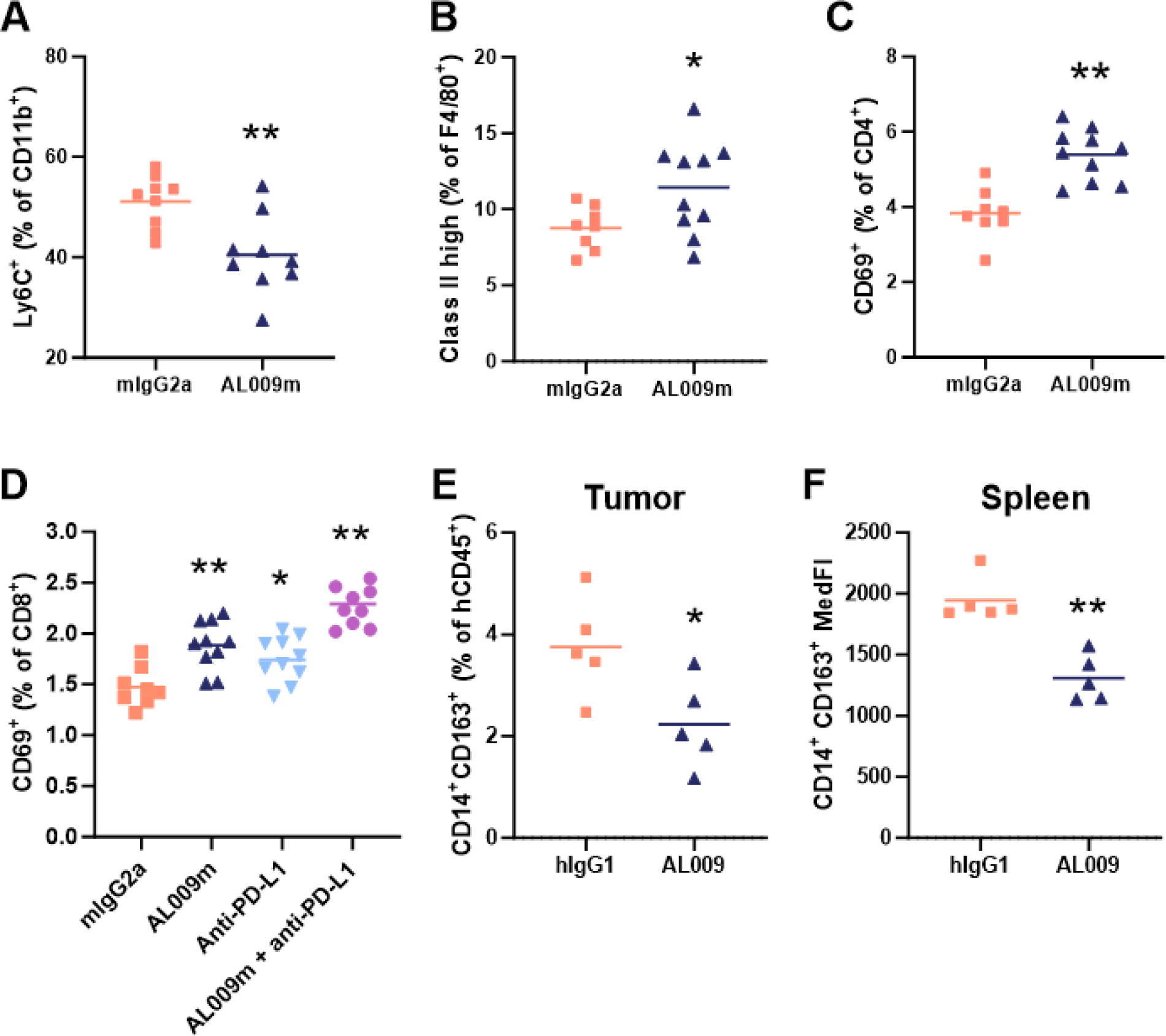
Pharmacodynamic biomarker data provides evidence that AL009m repolarizes myeloid cells and activates T cells in vivo. In the E0771 mouse tumor model, AL009m treatment (**A**) decreased MDSCs in tumor tissue as shown by a decrease in Ly6C^+^ cells, (**B**) activated macrophages in spleen tissue as shown by increased Class II high cells, and (**C**) activated CD4^+^ T cells, as shown by an increase in CD69^+^ cells. (**D**) CD8^+^ T cells were also increased with anti-PD-L1 treatment alone or in combination with AL009m treatment. (**E-F**) In a humanized mice tumor model, AL009 treatment decreased levels of the pharmacodynamic biomarker CD163, a marker for suppressive M2 macrophages, in the tumor and spleen. MDSC, myeloid-derived suppressor cell; MedFI, median fluorescence intensity. Mean is shown. \**p* < 0.05, \*\**p* < 0.01, 2-sided t-test.

## DISCUSSION

The current state of immunotherapy offers significant promise to treat cancer; however, while up to half of all patients may be eligible for treatment with immunotherapy, less than 15% derive clear benefit (25). TAMs heavily influence the TME and may comprise up to 50% of tumor mass, which creates barriers to anti-tumor immunity and may underlie the resistance of some patients to checkpoint inhibitors such as anti-PD1, anti-PD-L1, and anti-CTLA4 (26). Consequently, an aspect of cancer immunotherapy drug development has shifted focus to targeting the suppressive myeloid cells associated with tumors. There are currently TAM-targeting therapeutics under investigation which aim to either deplete TAMs, block recruitment of macrophages to the tumor, or reprogram TAMs to a proinflammatory phenotype (26, 27). Depletion and blocking recruitment of macrophages disregard the anti-tumor potential of creating reprogrammed, proinflammatory macrophages in the TME (27). Reprogramming TAMs to a proinflammatory phenotype results in stimulation of cytotoxic T cells and tumor regression in preclinical studies (26). Therapeutics aiming to reprogram TAMs may be effective as a monotherapy or in synergy with checkpoint inhibitors (26, 28). As a TAM-reprogramming therapeutic, AL009 provides two potential avenues for enabling immunotherapeutic activity: as a monotherapy to drive macrophage modulation of the TME towards a more immune-responsive environment, and in combination with checkpoint inhibitors to disrupt inherent immune-mediated inhibition of the adaptive immune response. AL009 therefore represents a novel, multi-functional therapeutic that has capacity for interrupting multiple negative regulatory signals in TAMs with significant potential for treating cancer.

With its Siglec-9 ECD and engineered Fc region, AL009 targets immune cells to widely block inhibitory signaling from Siglec receptors, leading to reprogramming of suppressive macrophages. Siglecs are responsible for regulating immune homeostasis, with inhibitory Siglecs playing a critical role in dampening immune responses (4, 8). The data from our experiments confirm the ability of the Siglec-9 ECD of AL009 to bind sialic acid ligands of not only Siglec-9 but also other inhibitory Siglecs, thus broadly blocking immune-inhibitory Siglec–sialic acid interactions. The disruption of this interaction allows anti-tumor activation signaling to proceed (Fig. 8). The engineered Fc portion of AL009 facilitates localization to immune cells and cooperative binding with the Siglec-9 ECD results in increased activity on innate immune cells. (Fig. 8).

**Fig. 8.**
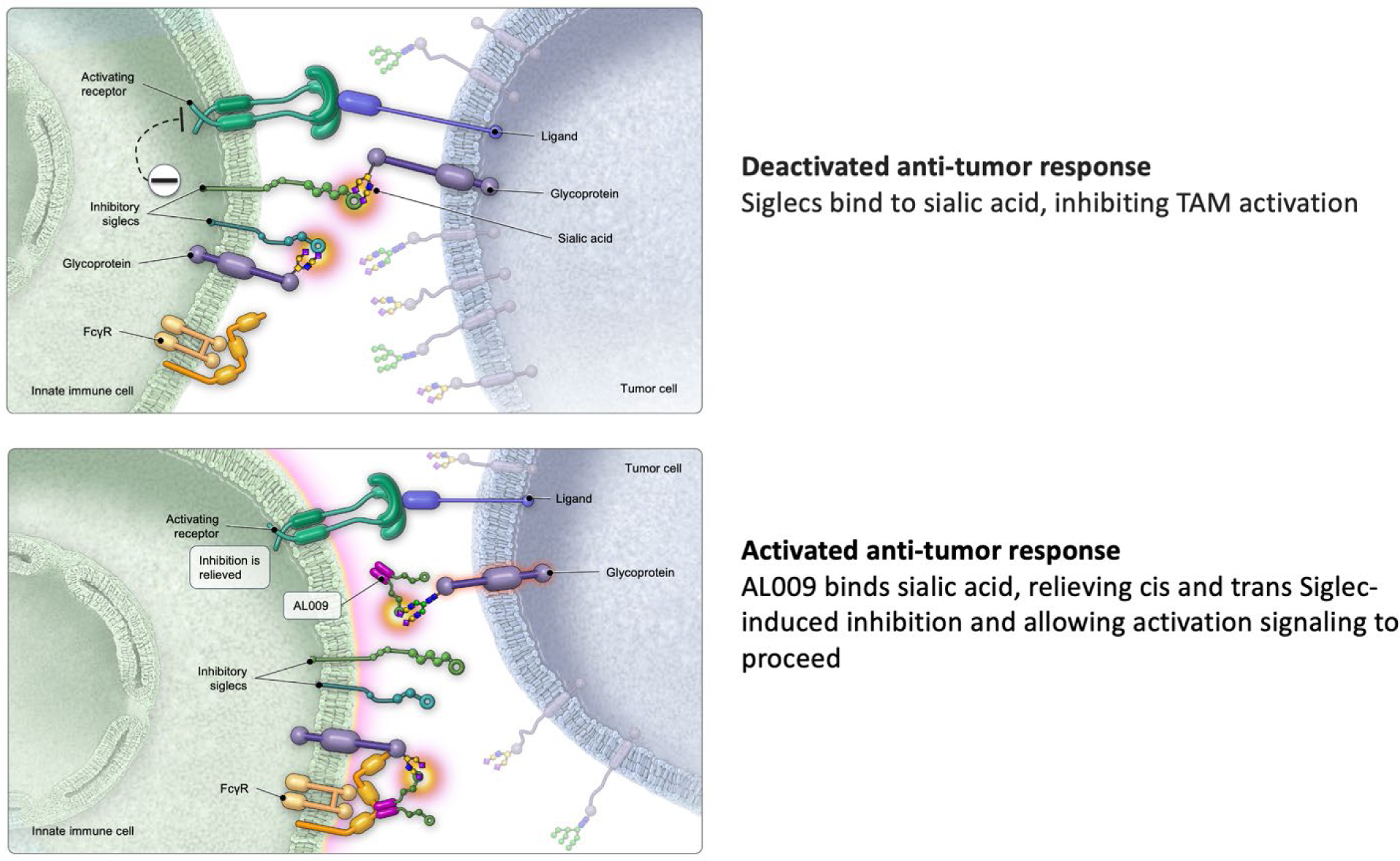
AL009 mechanism of action. AL009 consists of an extracellular Siglec domain and an engineered Fc portion for immune cell targeting. The Siglec portion of AL009 binds sialic acid ligands in a cis or trans fashion, relieving Siglec-mediated immune suppression. FcγR, Fcγ receptor.

Overall, AL009 is a differential approach to test the therapeutic hypothesis that TAM reprogramming is an effective method of generating an anti-tumor effect in humans. Thus far, pre-clinical studies have not identified any significant toxicology signals. In addition, serum and immune cell pharmacodynamic biomarkers have been identified, and immunohistochemistry companion diagnostic development is underway. Taken together, these preclinical results support clinical testing of the safety and efficacy of AL009.

## Acknowledgments

Editorial support and publication assistance were provided by SCIENT Healthcare Communications (Cedar Knolls, NJ, USA).

## Abbreviations used in this article

ADCC: antibody-dependent cellular cytotoxicity
ADCP: antibody-dependent cellular phagocytosis
ECD: extracellular domain
Fc: fragment crystallizable
FcγR: Fcγ receptor
IFNɣ: interferon gamma
MDSC: myeloid-derived suppressor cell
MedFI: median fluorescence intensity
Siglec: sialic acid–binding immunoglobulin-type lectin
TAM: tumor-associated macrophage
TME: tumor microenvironment
TREM-1: triggering receptor expressed on myeloid cells 1
TREM-2: triggering receptor expressed on myeloid cells 2

