## Supplemental Figures 2-4 for "The multi-Siglec inhibitor AL009 reprograms suppressive macrophages and activates innate and adaptive tumor immunity"

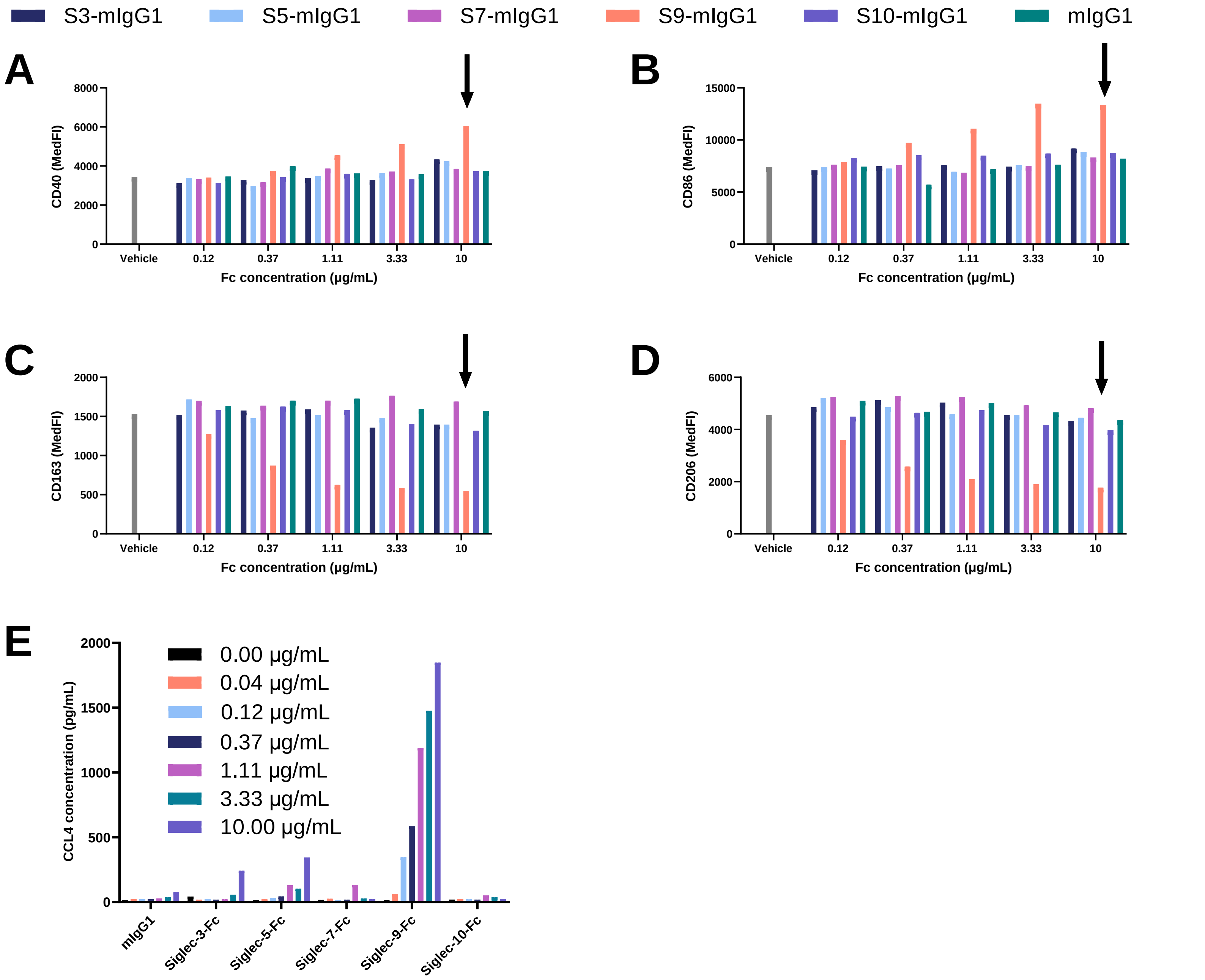


**Fig. S2**. Siglec-9-Fc has a unique impact on MDSC (**A-B**) activation and (**C-D**) suppression markers. (**E**) A dose-dependent increase in CCL4 production by MDSCs was observed with Siglec-9-Fc treatment. The ECDs of Siglec-3, -5, -7, -9, and -10 were expressed as mouse IgG1 fusion proteins and incubated with human MDSCs for 48 hours. Flow cytometry to assess cell surface molecules and analysis of conditioned media for chemokines were performed.

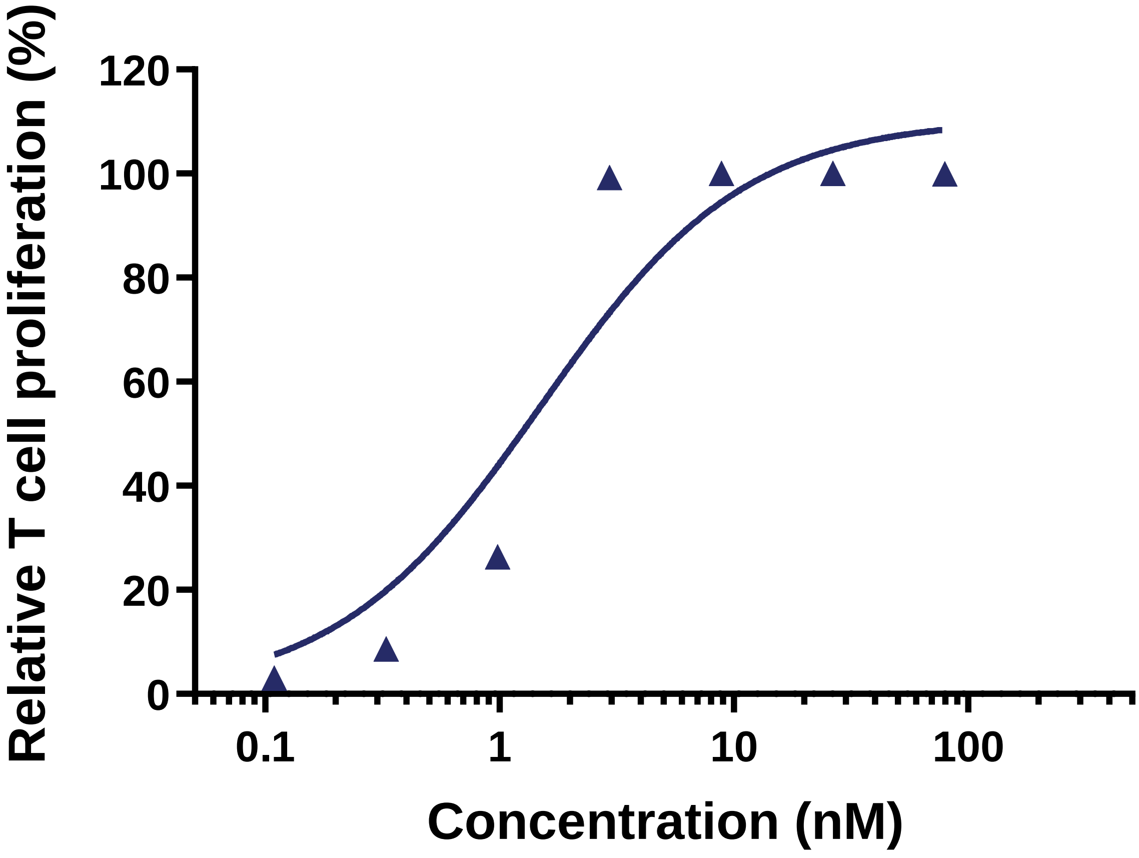


**Fig. S3**. AL009 relieved MDSC-mediated suppression of T cells with an $EC_{50}$ of 1-2 nM.

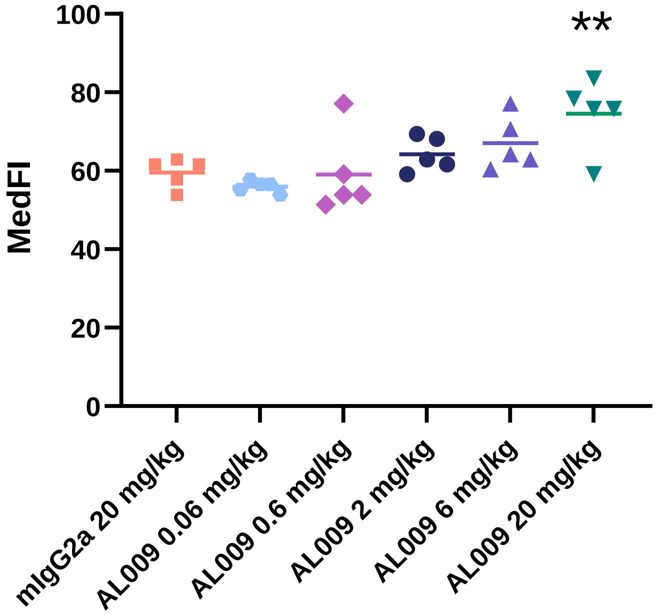


**Fig. S4**. AL009 induced dose-dependent increases in CD86 on monocytes. **p < 0.01, 2-sided t-test.
